# A Neural Model of RNA Splicing: Learning Motif Distances with Self-Attention and Toeplitz Max Pooling

**DOI:** 10.1101/2021.05.24.445518

**Authors:** Thomas P. Quinn, Dang Nguyen, Sunil Gupta, Svetha Venkatesh

## Abstract

Alternative RNA splicing is an important regulator of tissue development and specificity, and a relevant mechanism in cancer progression. There exists a strong motivation to disentangle the rules that govern RNA splicing, in part because this knowledge may one day yield new clinically relevant diagnostic tools and therapeutics. It is no easy to task to reverse engineer how the splicesome machinery choreographs the removal and addition of RNA elements following transcription. Here, we propose an interpretable neural network called the Toeplitz ATtention Architecture (TATA), which learns distance-dependent motif interactions through a novel Toeplitz max pool layer that captures the relative distance between interacting CNN filters. TATA is a completely transparent “clear-box” solution: every model parameter is human-interpretable. We validate TATA on simulated data, then apply it to real data to identify putative cis-regulatory elements that interact with primary RNA splice sites.

## 1 Introduction

Within a cell, DNA is transcribed as RNA which is eventually translated to make proteins. Before a protein is made, the RNA may be altered through post-transcriptional modification. One such modification, called *RNA splicing*, involves “cutting out” parts of the RNA and “glueing together” the remaining parts. In *alternative splicing*, a cell can turn a single RNA precursor into several different RNAs by changing which parts are “cut out”. Alternative splicing is an important regulator of tissue development and specificity [Baralle and Giudice, 2017], and a key driver of cancer progression [Bonnal et al., 2020]. Computational models of RNA splicing can help researchers discover novel splice mechanisms that may be targetable by therapeutic agents [Garcia-Blanco et al., 2004], *but interpretable models are needed to succeed in this endeavour*.

The convolutional neural network (CNN) is a natural choice for modelling RNA splicing. The first layer of a CNN meaningfully encodes the RNA through translation-invariant filters called *motifs* [Eraslan et al., 2019]. The deeper layers of a CNN then encode the nested interactions between the motifs. Deep CNNs perform well empirically (e.g., [Koo and Eddy, 2019, Wang et al., 2019, Jaganathan et al., 2019, Bao et al., 2019, Albaradei et al., 2020]), but are opaque [Rudin, 2019]: the neural connections do not readily provide a concise knowledge representation of the learned motif interactions. This opacity is intrinsic to the deep CNN itself, arising from a mismatch between the operations of the model and the nature of human-scale reasoning [Burrell, 2016]. There is a need for new interpretable models of RNA splicing, capable of learning a concise knowledge representation of motifs and their interactions.

RNA splicing depends on more than just motifs: it also depends on *context*, most notably the distance between cooperating motifs [Ule and Blencowe, 2019]. Quinn et al. [2020] define the *distance-dependent motif interaction problem* which states that a genomic event will occur if and only if (i) two (or more) motifs are present and (ii) the motifs are separated by a gap of a specific size (see Figure 1). Learning an interpretable solution to this problem is challenging. Although one recent study proposed an interpretable motif interaction model, it uses an expensive optimization procedure that makes a strong assumption about the distributional nature of intermotif distances [Quinn et al., 2020]. In contrast to [Koo et al., 2018, Avsec et al., 2020], who use post-hoc explainable AI (XAI), *we seek to describe distance-dependence directly through model parameters, whereby interpretability is built-in a priori as an intentional design choice*.

**Figure 1:**
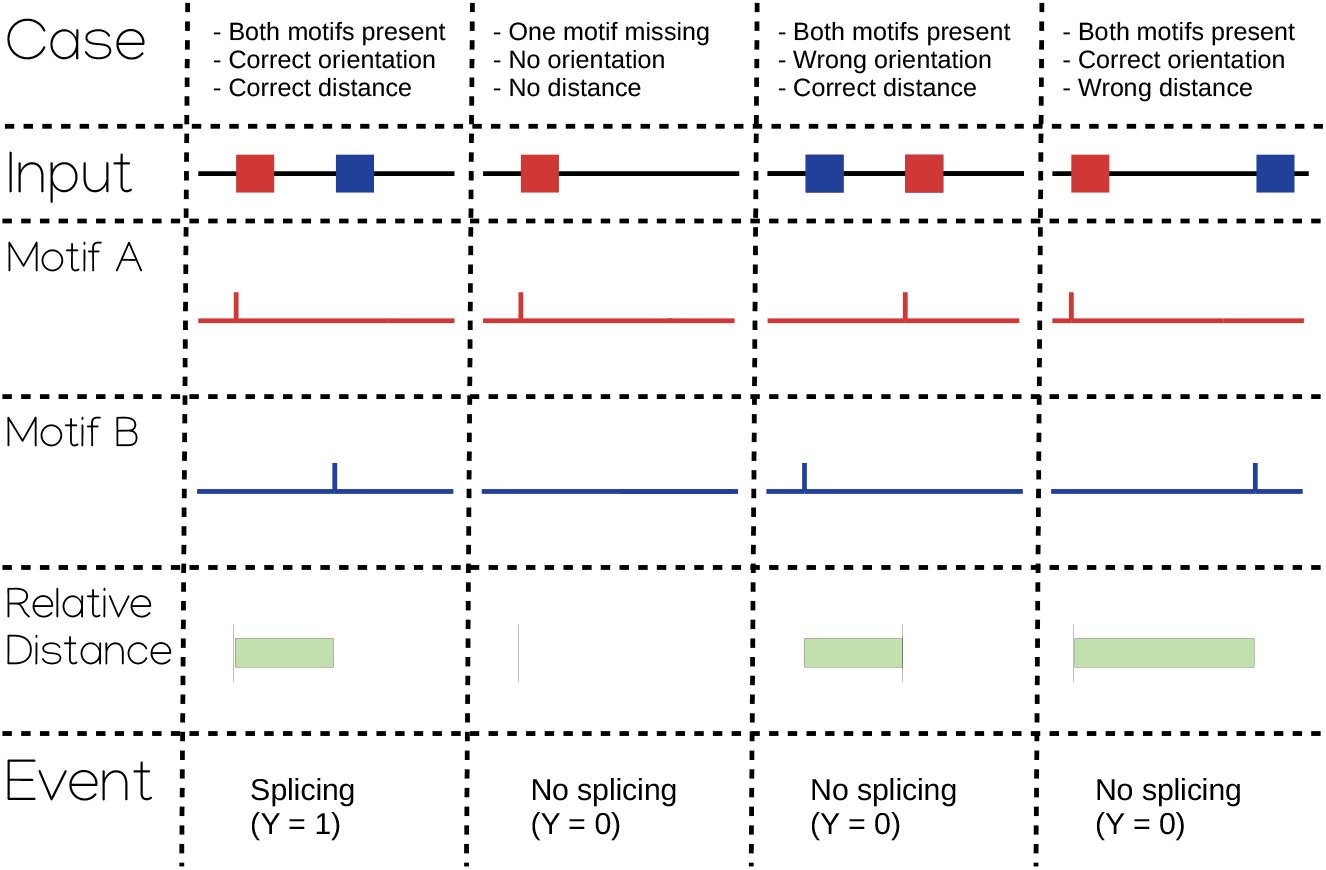
A visual summary of the distance-dependent motif interaction problem. In the first column, two motifs are separated by the correct distance and therefore *y* =1. In the second column, one motif is missing and therefore *y* = 0. In the third column, two motifs are separated by the correct distance but have the wrong orientation and therefore *y* = 0. In the fourth column, two motifs are separated by the incorrect distance and therefore *y* = 0. An interpretable model should explicitly identify the relevant motifs and their distance dependence.

We propose the Toeplitz ATtention Architecture (TATA) as an explicitly interpretable solution to the distance-dependent motif interaction problem. TATA adapts self-attention to learn (i) the identity of the relevant motifs and (ii) the importance of the relative distances between them (including orientation). Furthermore, it advances RNA modelling along 2 dimensions:

1. **TATA does not assume an inter-motif distance distribution:** Our novel Toeplitz max pool layer sits on top of a self-attention head, encoding the relative distance between motifs as an interpretable hidden layer. This requires no prior knowledge about the distribution of the relative distances.
2. **TATA can learn interactions between discrete sets of motifs:** A continuous relaxation of motif set membership allows the neural network to learn discrete motif sets, which in turn allows the neural network to learn the interaction between sets, thus capturing more complex motif interactions.

TATA learns distance-dependent motif interactions through a novel Toeplitz max pool layer, built on top of a modified self-attention head, that captures the relative distance between interacting CNN filters. TATA is a completely transparent “clear-box” solution: every model parameter is human-interpretable. We validate TATA on simulated data, then apply it to real data to identify putative cis-regulatory elements that interact with primary RNA splice sites.

## 2 Related Background

### Biological motifs

A DNA/RNA sub-sequence is biologically relevant if it is recognized at a molecular level. Different sub-sequences can cause the same genomic event. A motif is an abstraction of multiple related sub-sequences. Thus, a motif is more like a regular expression than an ASCII string. For example, the U1 snRNP protein could initiate RNA splicing by binding to any sub-sequence captured by the regular expression ‘(C|A)AGGU(G|A)AG’ [Mount et al., 1992]. Motif activity may depend on the relative distance between interacting motifs [Ule and Blencowe, 2019], for example in the case of multivalent binding by RNA-binding proteins [Sohrabi-Jahromi and Söding, 2021]. Often, there is no “perfect” inter-motif distance; rather, there exists a range of valid inter-motif distances. For example, in RNA splicing, the inter-motif distance between the branch point and the 3’ consensus splice site is usually 16-to-25 bases apart [Ohno et al., 2018].

### CNNs for motif discovery

The idea of using computer algorithms to learn biological motifs is not new. For example, Stormo et al. [1982] learned a weight matrix called a *position weight matrix* that could predict whether a gene is valid. These weights, when normalized, represent the expected relative importance of each DNA base at each position, and thus define a *consensus motif*. Recently, deep learning methods have been used for motif discovery. Among model-agnostic approaches, Zhou and Troyanskaya [2015] proposed *in silico mutagenesis*, a technique that perturbs the DNA input and measures how the perturbation changes prediction. Among model-specific approaches, CNNs are a popular choice because the first layer kernel weights represent a position weight matrix. Studies have looked at how exponential activation [Koo and Ploenzke, 2019], Gaussian noise injection [Koo et al., 2019], and kernel regularization and constraints [Ploenzke and Irizarry, 2018] could make the resultant position weight matrices more robust. In TATA, we also use a convolutional layer to learn consensus motifs.

### CNNs for motif interactions

Jaganathan et al. [2019] proposed a 32-layer deep CNN with 696,739 learnable parameters to predict splice site presence from DNA input. The high accuracy of this model suggests that motif context, including inter-motif distance, can be learned implicitly by the hidden layers of a CNN. We know of two studies that infer motif context rules from a deep CNN, though neither studied RNA splcmotifs from Stage 1 are grouped iing directly. Koo et al. [2018] and Avsec et al. [2020] both take a second-order *in silico mutagenesis* approach where one motif is fixed while another is inserted systematically at different relative distances to give a model-agnostic explanation of distance-dependent interactions. Although their implementations are intuitive, model-agnostic explanations can be unreliable: *an explanation of how a model appears to work may not be how the model actually works* (c.f., [Alvarez-Melis and Jaakkola, 2018a, Rudin, 2019]). In studying RNA splicing, Quinn et al. [2020] propose an explicitly interpretable model of inter-motif distance, but use an expensive nested optimization procedure, and restrict their analysis to pairwise motif interactions. In TATA, we also propose an explicitly interpretable model of inter-motif distance, but remove the distributional assumption and further capture interactions between motif sets.

### Attention for biological sequences

In the quest for understanding how a model works, researchers have proposed methods for *probing* the black-box [Azodi et al., 2020]. Notable examples include methods that assign *saliency*, a measure of feature importance, to input variables based on gradients [Simonyan et al., 2014]. By selecting a reference input, importance scores can be computed for individual samples [Shrikumar et al., 2019]. In contrast, [Bahdanau et al., 2016] uses the attention mechanism to allow a model to automatically search for the important parts of an input. Attention, being related to the concept of *self-explanation* [Alvarez-Melis and Jaakkola, 2018b], can assign saliency from within the model, and has been previously used in genomics research to produce interpetable sample-specific importance scores [Beykikhoshk et al., 2020]. Attention is now an important part of transformer architectures [Vaswani et al., 2017], a light-weight alternative to CNNs for big data like whole genomes [Clauwaert and Waegeman, 2020]. Within transformers are *self-attention heads* that compute the attention between different positions of the same input. In TATA, we adapt self-attention to learn distance-dependent motif interactions.

## 3 Model Architecture

### 3.1 Overview

TATA uses self-attention to solve the distance-dependent motif interaction problem. It has 4 stages that learn to represent (1) motif discovery, (2) their interactions, (3) their context dependence, and (4) their contribution to the final prediction (see Figure 2).

- **Stage 1: Motif discovery** A convolutional layer learns to represent motifs, where the weights from one filter correspond to one motif.
- **Stage 2: Motif interaction** The motifs from Stage 1 are grouped into sets, then selfattention is used to capture multiplicative interactions along the RNA sequence.
- **Stage 3: Context dependence** The self-attention head is summarized to represent the relative distance between interacting motifs.
- **Stage 4: Final regression** A generalized linear regression makes a final prediction.

**Figure 2:**
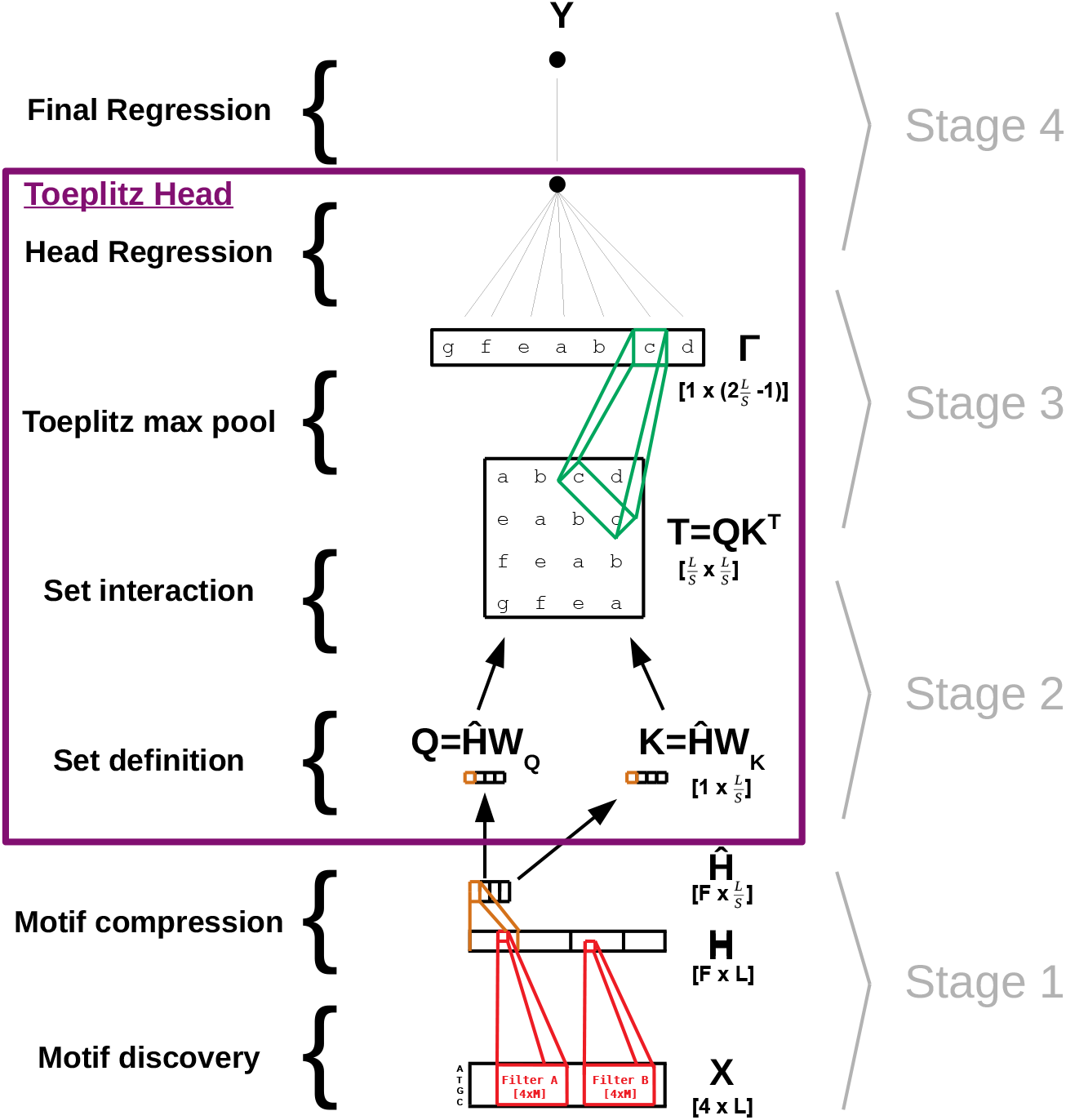
An overview of the TATA model, showing a single *Toeplitz head*. Stage 1 discovers the identity of the motifs and then groups them into motif sets (see Section 3.2). Stage 2 captures the interactions between motif sets (see Section 3.3). Stage 3 assigns weight to a motif set interaction based on the relative distance between the sets (see Section 3.4). Stage 4 makes the final prediction (see Section 3.5). All parameters are explicitly interpretable. *K* Toeplitz heads can be stacked in parallel to learn *K* independent interacting sets.

### 3.2 Stage 1: Motif discovery

Let us consider a total of *N* DNA sequences **X** with *p* bases having 4 possible states, 1-hot encoded as a [*N* × 4 × *L*] tensor (the 4 states being adenine, cytosine, guanine, or thymine). We pass the input through a convolutional layer having *F* filters of size [*M* × 4], where *F* is the number of motifs and *M* is the maximum motif length:

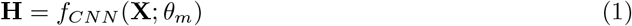

where the parameters *θ_m_* represent the motif identities (as visualized after normalization by a softmax transform). The resultant **H** has the dimension [*N* × *F* × *L*] and describes motif presence at each position along the DNA sequence.

We then pass **H** through a max pool layer to summarize motif presence in “chunks”. This motif compression step reduces the resolution at which the architecture detects motif interactions, but advantageously decreases the number of downstream parameters and the runtime of each epoch.

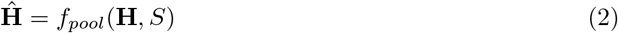

where *S* is both the kernel size and the stride. The resultant **Ĥ** has the dimension 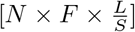 (or rounded up in the case that *S* is not a multiple of *L*).

### 3.3 Stage 2: Motif interaction

Since motifs carry meaning to domain experts in the field of genomics, **H** is an interpretable embedding of the DNA sequence. One could pass this embedding through a self-attention head, as done in natural language processing [Bahdanau et al., 2016].

To introduce self-attention into the model, we will use a variant of the attention head found in the transformer architecture (c.f., [Vaswani et al., 2017]). In transformers, the matrices **Q** and **K** are computed from the embedding matrix **Ĥ** as

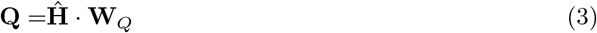

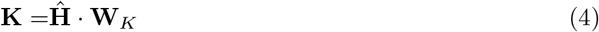

by the weights {**W**_*Q*_, **W**_*K*_}, each having the size [*F* × *D*]. We interpret **Q** (and **K**) to represent the presence of the motif sets. The weights **W**_*Q*_ and **W**_*K*_ define the sets, where a positive entry at coordinate {*F*, *D*} indicates that motif *F* belongs to set *D*. A non-negative weights constraint ensures **W**_*Q*_ ≥ 0 (and **W**_*K*_ ≥ 0), reinforcing this intuition. Although these weights are not discrete, we can make them discrete with an algorithm described in the Implementation section.

The transformer attention head eventually uses 3 matrices to compute a latent representation of the attended input. In TATA, we are only interested in the query-key component

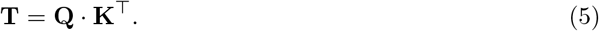

The matrix **T** has the size 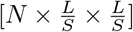, *and encodes the multiplicative interactions between motif sets*. The term “interaction” follows from the statistical literature on non-additive effects [Tsang et al., 2018]. In **T**, all possible pairwise multiplicative interactions are calculated at once by taking the dot product between the **Q** and **K** embeddings.

### 3.4 Stage 3: Context dependence

Each coordinate {*i*,*j*} in **T** represents the magnitude of the interaction between the two sets at *j* — *i* positions apart.^1^ Due to the structure of the matrix, all elements of a given off-diagonal represent the same relative distance. For example, consider the [4 × 4] Toeplitz matrix

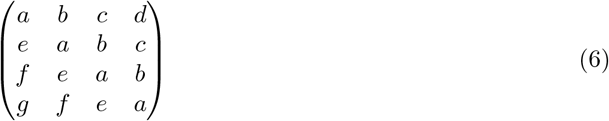

Here, the off-diagonal *b* would represent all interactions at 1 position apart, *c* at 2 positions apart, and *d* at 3 positions apart (while *e*, *f*, and *g* represents the opposite orientation).

Our objective is to represent distance-dependent interactions *explicitly*. One way to do this is to take the maximum for each off-diagonal. This gives us a measure of the strongest interaction for a given relative distance. Put symbolically, we compute the maximum value *γ_d_* for each integer 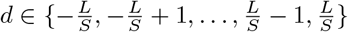

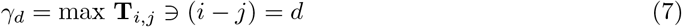

where each *d* represents one fixed inter-motif distance (e.g., *d* = 2 captures motif set interactions that are 2 bases apart). We then concatenate these as:

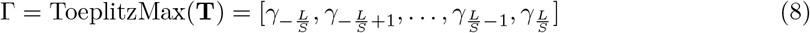

We call this operation the *Toeplitz max pool* because it reduces the matrix into a vector that describes a diagonal-constant matrix of the maximum off-diagonals. Thus, Toeplitz max pooling reduces the 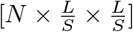 interaction matrix to a 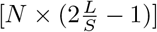 vector.

We feed forward the results from the Toeplitz max pool layer to a single node e which represents the *evidence* provided by a single self-attention head. Although we could connect the relative distance vector Γ to e using any differentiable function, we simplify the interpretation by using a direct connection with linear activation so that

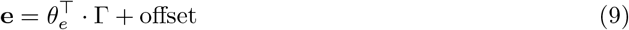

where the weights *θ_e_* represent the importance of each off-diagonal.

### 3.5 Stage 4: Final regression

For *k* ∈ {1…*K*} attention heads stacked in parallel, each attention head will produce its own piece of evidence *e_(k)_*. We can concatenate these to form the penultimate layer

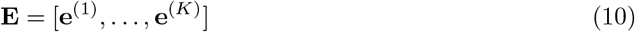

which we use to estimate the final output **Y** as **Ŷ** via a generalized linear model

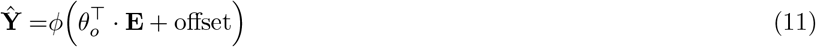

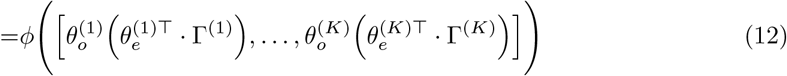

where the weights *θ_o_* represent the importance of each attention head, and *φ* is the link function chosen based on the nature of **Y** (e.g., continuous or binary). The parameters are then learned by minimizing the loss between **Y** and **Ŷ**.

## 4 Implementation

### 4.1 Initialization by knowledge transfer from Deep CNN

Learning the motifs and relative distances simultaneously presents a difficult optimization problem (c.f., [Quinn et al., 2020]). Although we could train TATA end-to-end with back-propaganation, the vanilla architecture has variable performance depending on the initialization of the motif identity layer weights *θ_m_*. We bring greater stability to the training regime by transferring learned motif identities from a Deep CNN whose first layer is identical to TATA’s first layer. Since the first layers are identical, the weights can be transferred by direct copy.

Once transferred, the weights *θ_m_* are frozen and TATA is trained end-to-end to learn **W**_*Q*_, **W**_*K*_, *θ_e_*, and *θ_o_*. After convergence, the weights *θ_m_* are unfrozen and TATA is trained again to fine-tune all weights. The exact Deep CNN architecture is described in Experiments.

### 4.2 Set hardening

We interpret the weights **W**_*Q*_ and **W**_*K*_ to describe a set of motifs that interact together. Although we use a non-negative weights constraint to force positive values, the weights are otherwise free to take on any continuous value. As such, the sets described by **W**_*Q*_ and **W**_*K*_ are not really discrete sets *per se*. This complicates their interpretation.

We use a *weight hardening procedure* to discretize the weights. In weight hardening, the continuous weights **W** are replaced with discrete weights 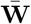 based on a cutoff *c*

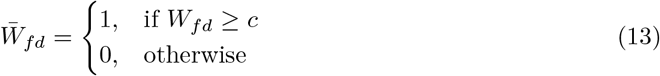

for *f* ∈ {1…*F*} filters and *d* ∈ {1…*D*} sets. It follows that when the cutoff *c* is very large, the matrices **W**_*Q*_ and **W**_*K*_ will become very sparse. As such, set hardening also acts as a kind of regularization [Gordon-Rodriguez et al., 2021].

By conceptualizing the weights as a *continuous relaxation* of the ideal discrete weights [Maddison et al., 2017], we can use the training data to identify an *optimal cutoff c*\*, where replacing **W** with 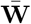 causes the smallest decrease in accuracy. To find the optimal cutoff, we trial *C candidate cutoffs* using the *C* largest unique values in {**W**_*Q*_, **W**_*K*_}. For each candidate cutoff, we discretize the weights and recompute the downstream layer activations.

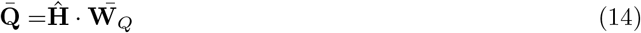

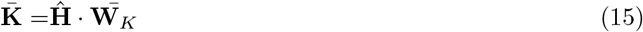

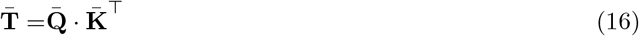

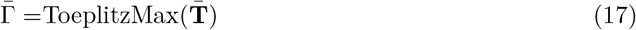

Then, for each cutoff, we use the training data to fit a “new model” to predict the outcome from 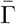. Over 5 folds, we compute an average AUC and standard error, and select the largest cutoff (i.e., the most sparse model) whose highest AUC score is above the *maximum average AUC, minus 1 of its standard error*. Here, the “new model” is a random forest (RF), implemented using the randomForest R package with default arguments except `ntrees=25`. Once the optimal cutoff is chosen, the continuous weights are made discrete and frozen; then, the model is fine-tuned.

### 4.3 Training

The final loss is binary cross-entropy without any regularization. TATA is implemented in keras, and trained using the ADAM optimizer with a learning rate of 0.001. Most hyper-parameters are chosen *a priori* based on prior domain knowledge or intuition. Table 1 describes all relevant hyper-parameters, and the justification for their set value.

**Table 1:**
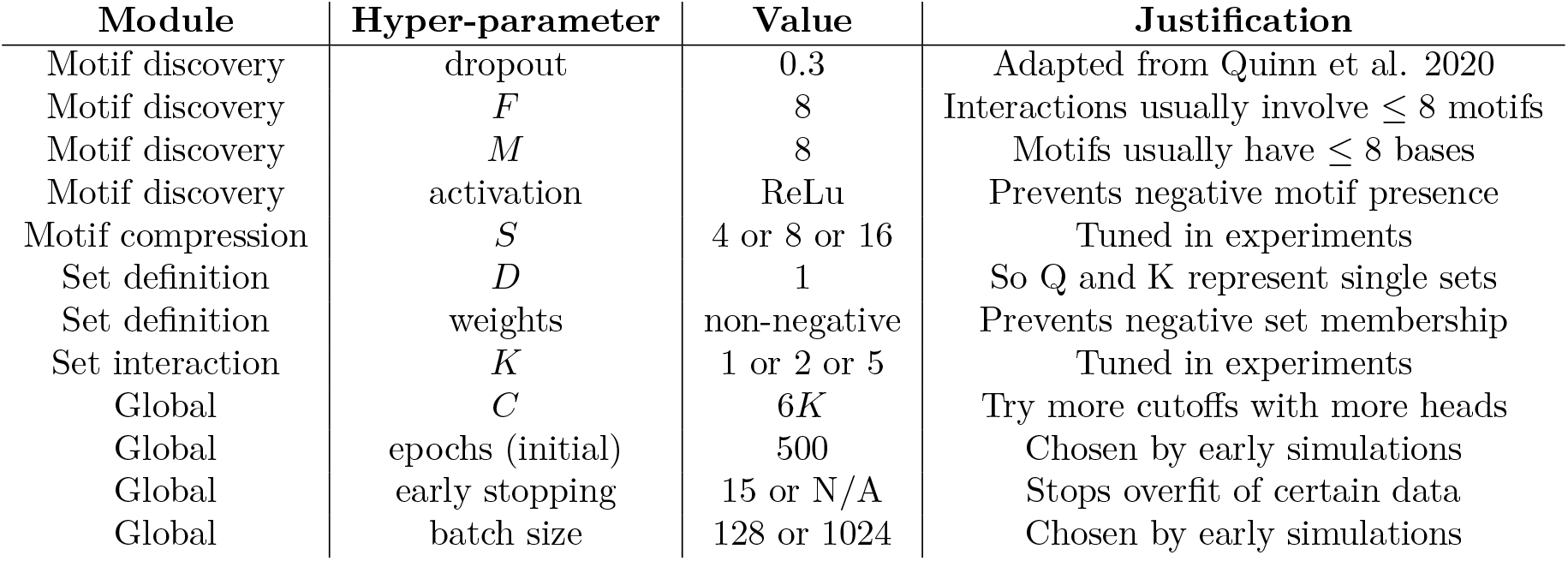
The hyper-parameter choices for our TATA architecture.

| Module | Hyper-parameter | Value | Justification |
| --- | --- | --- | --- |
| Motif discovery | dropout | 0.3 | Adapted from Quinn et al. 2020 |
| Motif discovery | $F$ | 8 | Interactions usually involve $\leq 8$ motifs |
| Motif discovery | $M$ | 8 | Motifs usually have $\leq 8$ bases |
| Motif discovery | activation | ReLU | Prevents negative motif presence |
| Motif compression | $S$ | 4 or 8 or 16 | Tuned in experiments |
| Set definition | $D$ | 1 | So Q and K represent single sets |
| Set definition | weights | non-negative | Prevents negative set membership |
| Set interaction | $K$ | 1 or 2 or 5 | Tuned in experiments |
| Global | $C$ | $6K$ | Try more cutoffs with more heads |
| Global | epochs (initial) | 500 | Chosen by early simulations |
| Global | early stopping | 15 or N/A | Stops overfit of certain data |
| Global | batch size | 128 or 1024 | Chosen by early simulations |

## 5 Experiments

### 5.1 Data

We demonstrate TATA on 2 series of simulated data sets (with 3 data sets per series), 2 real-world publicly available data sets (acceptor and donor), and 2 hypothesis-driven subsets of the real-world data. For each data set, we measure AUC on 5 independent training-test set splits (see Table 2).

- **Simulated Series 1:** These simulations represent *2-way Motif Interactions*. They are nucleotide FASTA files simulated to be an intron sequence, followed by a branch point (BP), more intron sequence, a 3’ RNA splice site (3SS), then finally by an exon sequence. When *y* = 1, the BP is separated from the 3SS by an average of 21 bases (standard deviation 2). When *y* = 0, the BP is separated from the 3SS by an average of either 29, 37, or 45 bases (standard deviation 2). These correspond to *relative gaps* of 8, 16, or 24 bases. These gaps correspond to Hard, Medium, or Easy problems, respectively.
- **Simulated Series 2:** These simulations represent *3-way Motif Interactions*. They are nucelotide FASTA files simulated to be an intron sequence, followed by a branch point (BP), more intron sequence, a 3’ RNA splice site (3SS), then finally by an exon sequence. This series differs from the first series in that, when *y* = 1, the intron also contains a polypyrimidine tract (PPT). When *y* = 0, the sequence either (a) does not have a PPT, (b) does not have a 3SS, or (c) has a BP that is separated from the 3SS by a relative gap of 8, 16, or 24 bases. These gaps correspond to Hard, Medium, or Easy problems, respectively.
- **Acceptor and Donor:** These data are nucleotide FASTA files of (a) acceptor vs. non-acceptor sites or (b) donor vs. non-donor sites. The data originally come from Albaradei et al. [2020], as separated into 5 training and test sets by Quinn et al. [2020].
- **Acceptor and Donor subsets:** These data are derived from Albaradei et al. [2020] to study whether and how context-dependence predicts splicing among select non-canonical motifs. Most acceptor sites have the trinucleotide CAG. The first subset includes acceptor (and non-acceptor) sites with a TAG instead. Similarly, most donor sites have the trinucleotide GGT. The second subset includes donor (and non-donor) sites with an AGT instead. These nonrepresentative cases, called non-canonical sites, are chosen because the literature suggests that they may more likely depend on motif interactions.

**Table 2:** The dimensions of the data sets used to train our TATA architecture.

| Data Set | Max Length (L) | Total Folds (N) | Training (N) | Test (N) |
| --- | --- | --- | --- | --- |
| 2x Interactions | 161 | 15 | 2744 | 1352 |
| 3x Interactions | 159 | 15 | 2744 | 1352 |
| Acceptor | 102 | 5 | 38461 | 18944 |
| TAG subset | 102 | 5 | 8685 | 4277 |
| Donor | 102 | 5 | 40358 | 19878 |
| AGT subset | 102 | 5 | 9000 | 4433 |

### 5.2 Baselines

We consider 9 baselines:

- **NLP-like methods:** For comparison with Quinn et al. [2020], we ran all baselines according to their specifications. These baselines are natural language processing (NLP) methods. They are bag-of-words (BOW), term frequency-inverse document frequency (TFIDF) [Arora et al., 2018], and Sqn2Vec, a neural embedding method [Nguyen et al., 2019].
- **Shallow CNN:** This CNN resembles the first layers of TATA: Input → 30% Dropout → CNN (8 motifs, sized [8 × 4]) → Global Max Pool. TATA also uses 8 motifs sized [8 × 4].
- **Wide CNN:** This CNN also resembles the first layers of TATA, but has more parameters: Input → 30% Dropout → CNN (32 motifs, sized [32 × 4]) → Global Max Pool.
- **Wider CNN:** This CNN also resembles the first layers of TATA, but has more parameters: Input → 30% Dropout → CNN (64 motifs, sized [64 × 4]) → Global Max Pool.
- **Deep CNN:** The first layers of the Deep CNN resemble the first layers of TATA: Input → 30% Dropout → CNN (8 motifs, sized [8 × 4]). The following layers are: Strided Max Pool (sized [4 × 1]) → 30% Dropout → CNN (8 motifs sized [8 × 1]) → Strided Max Pool^2^ (sized [4 × 1]) → Output. This is also the network used to initialize TATA as described in the Implementation section.

The similarities between TATA and the CNNs make it possible to evaluate the contribution of the Toeplitz Max Pool module as compared with having zero or more hidden layers.

## 6 Results

### 6.1 TATA distills knowledge about motif interactions

TATA has 5 sets of weights. They each represent different aspects of RNA splicing. The constraints placed on these weights, along with the operations that relate them to the output, ensure that the weights are biologically relevant and human-interpretable:

- {*θ_m_*} from Eq 1 define the motifs themselves. A softmax transform normalizes the weights as a position weight matrix, which can be visualized via a seqLogo plot.
- {**W**_*Q*_, **W**_*K*_} from Eq 4 define the motif membership within the interacting sets. They are, by design, non-negative, and can be discretized through our *weight hardening procedure*.
- {*θ_e_*,*θ_o_*} from Eq 12 define the importance of each relative distance in predicting the outcome **y**. Although *θ_e_* and *θ_o_* are separate parameters within the model, they can be multiplied together to get a single importance score (see Eq. 12).

There are no additional parameters within the model. Thus, these parameters together provide a complete description of model behavior.

### 6.2 TATA learns 2- and 3-way interactions

Figure 3 shows the performance of the TATA architecture compared with baselines run on 6 simulated data sets. For the 2-way interactions, only the Deep CNN and TATA models achieve high accuracy. Here, TATA performs best, perhaps because the Toeplitz head can explicitly represent the exact mechanism we simulated, yielding a more generalizable prediction than the Deep CNN layer. For the 3-way interactions, TATA matches the Shallow, Wide, and Deep CNNs. The 3-way interactions may be easier for the CNN to model because many examples in the negative class have 1 of the 3 motifs missing.

**Figure 3:**
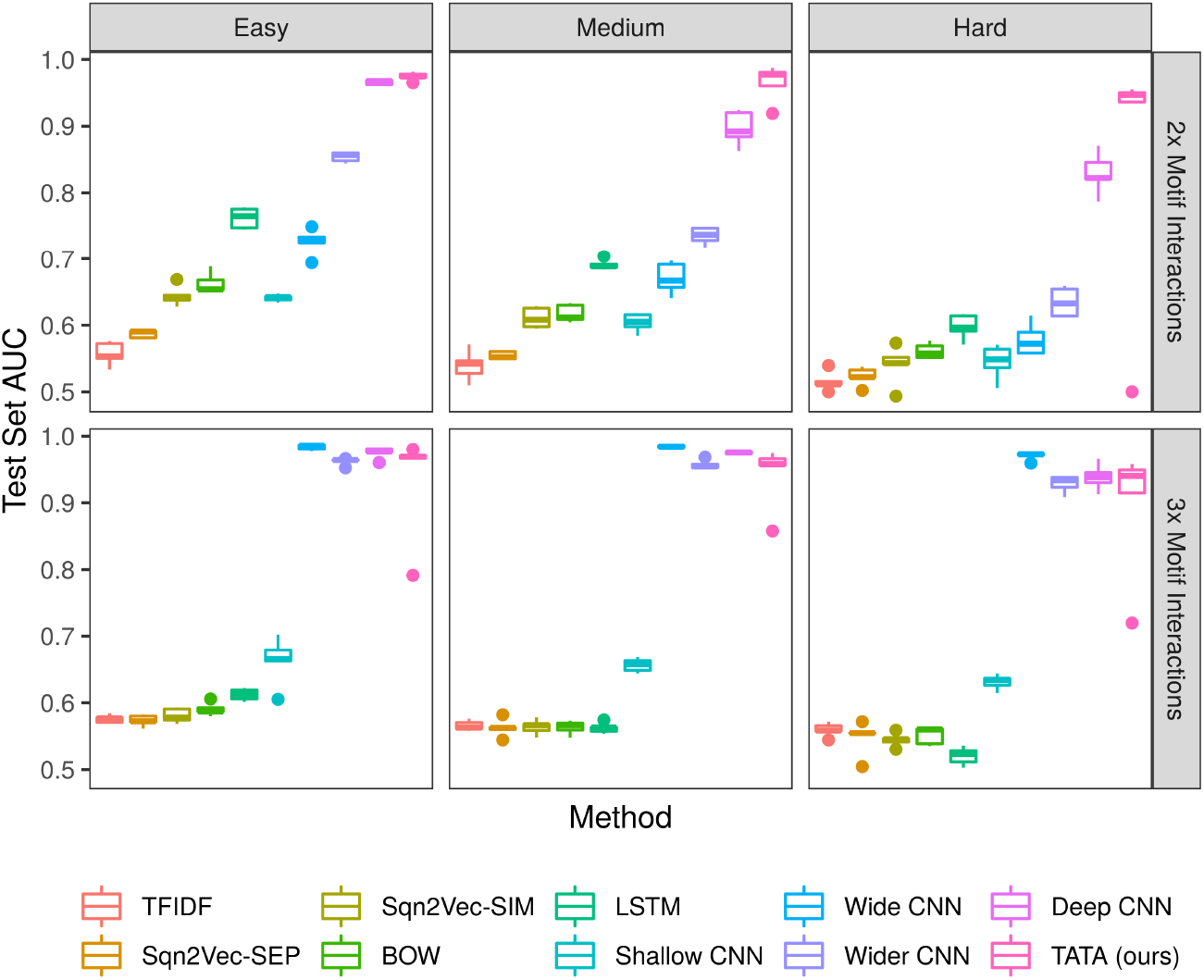
Test set AUC across 6 simulated data sets, using 5 independent training sets each. Our TATA architecture performs as well as, or better than, several state-of-the-art baselines. However, its parameters are explicitly interpretable. Here, we show TATA when *S* = 4 and *K* = 5. Hard, Medium, and Easy correspond to relative gaps of 8, 16, or 24 bases.

### 6.3 Continuous relaxation enhances mechanistic interpretations

The weights {**W**_*Q*_, **W**_*K*_} define motif set membership. To solidify the interpretation of {**W**_*Q*_, **W**_*K*_} as sets, we use a *weight hardening procedure* to discretize these parameters as 0 or 1. Figure 4 shows the relative difference in test set AUC before and after hardening, showing that hardening does not considerably impair model performance. Visually, we see that AUC drops by about 2% AUC on average, with a few outliers having worse performance.

**Figure 4:**
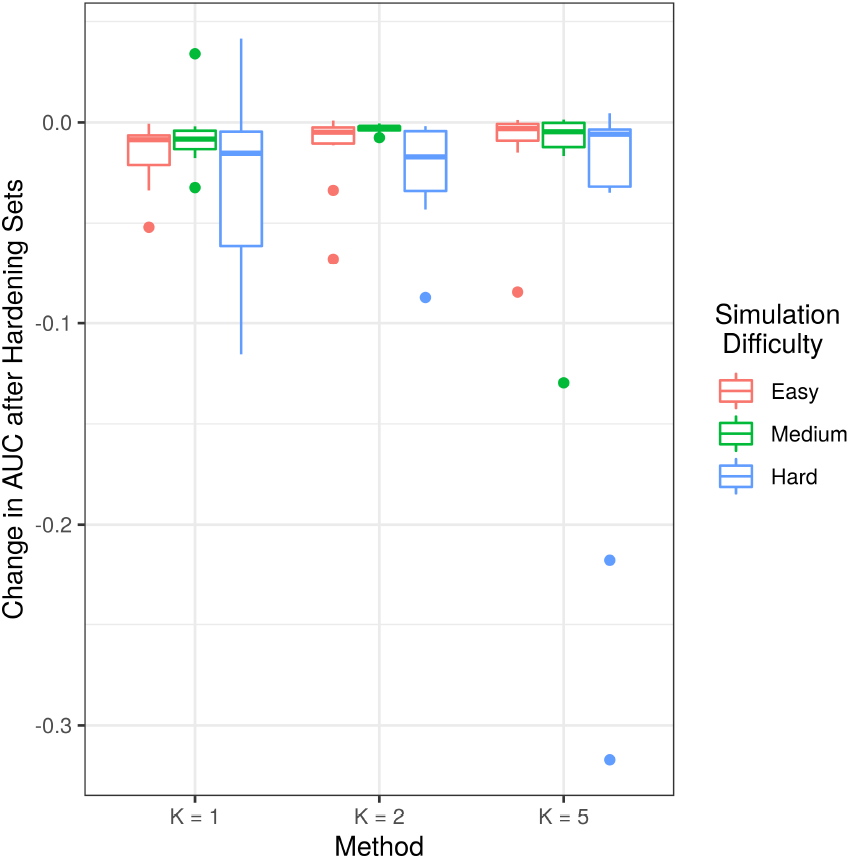
Change in test set AUC, before hardening vs. after hardening, across 6 simulated data sets, using 5 independent training sets each. Set hardening does not considerably impair performance, but can substantially improve interpretability.

With discretized sets, it becomes possible to describe each Toeplitz Max Pool head using a concise *visual-semantic representation*. Figure 5 provides one example, where every model parameter is combined into a single “dashboard” description of model behavior. Motifs, shown with seqLogo plots, are bracketed into sets. In this case, one set contains a single motif whose elaborate identity is better expressed as multiple distinct position weight matrices, perhaps as a compensation for motif compression. The line plot shows the importance of each relative distance, centered at a relative distance of 0. This figure, learned on simulated data, perfectly describes the rules used to simulate the data: the genomic event is most likely to occur when the branchpoint sequence (BPS, with regex ‘(C|T)T*A(C|T)’) is separated from the 3’ splice site (3SS, with regex ‘*(C|T)AGG’) by ~21 bases, and it is least likely to occur when the motifs are separated by ~45 bases.

**Figure 5:**
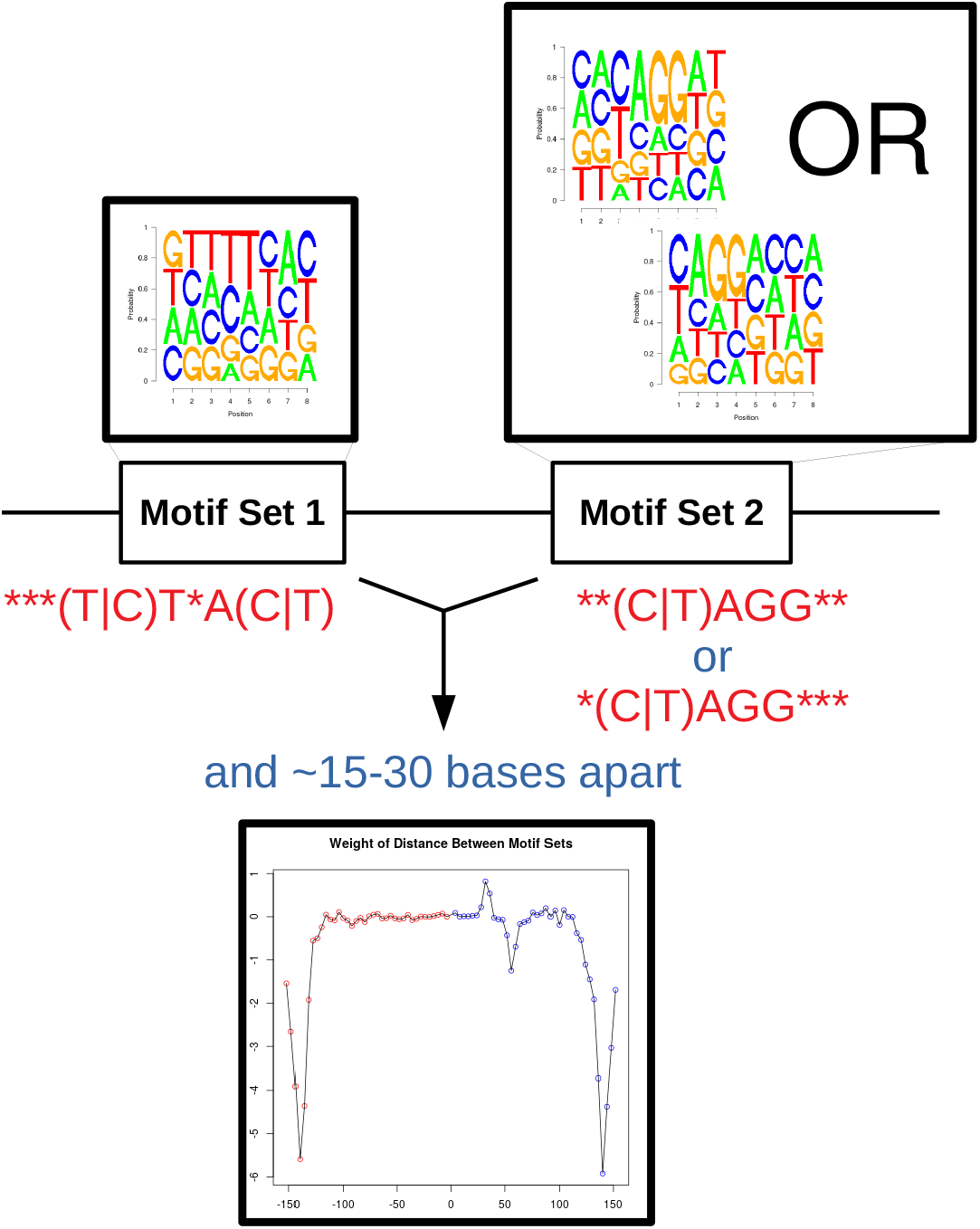
A concise visual-semantic representation of one Toeplitz Max Pool Head and all associated parameters. This representation, learned from the simulated data, accurately captures the ground-truth mechanism. The motifs, shown as seqLogo plots and regular expressions, are bracketed into sets that interact. The line plot shows the importance of the interaction for each relative distance. Here *K* = 1, but there would be *K* representations for *K* Toeplitz heads.

We note that high negative weight is also assigned when the BPS and 3SS are separated by ~ 150 bases in either direction. This appears to be an artefact of the simulation, whereby the positive class sequences tend to be shorter than the negative class sequences. As such, when the BPS and something 3SS-like are very far apart, for example by chance, the sequence must be very long and therefore assigned to the negative class. Such examples seem to appear rarely enough that the ground-truth mechanism is still detected.

### 6.4 Application to RNA Splice Site Prediction

Having established the reliability of the interpretations using simulated data, we benchmark TATA performance on real-world data, then carve out a hypothesis-driven subset of the data for further exploration. Figure 6 shows the average test set AUC for the 2 data sets and their subsets. Visually, we see that the TATA architecture is largely comparable with Shallow, Wide, and Deep CNNs, but with the added advantage of its parameters being explicitly interpretable.

**Figure 6:**
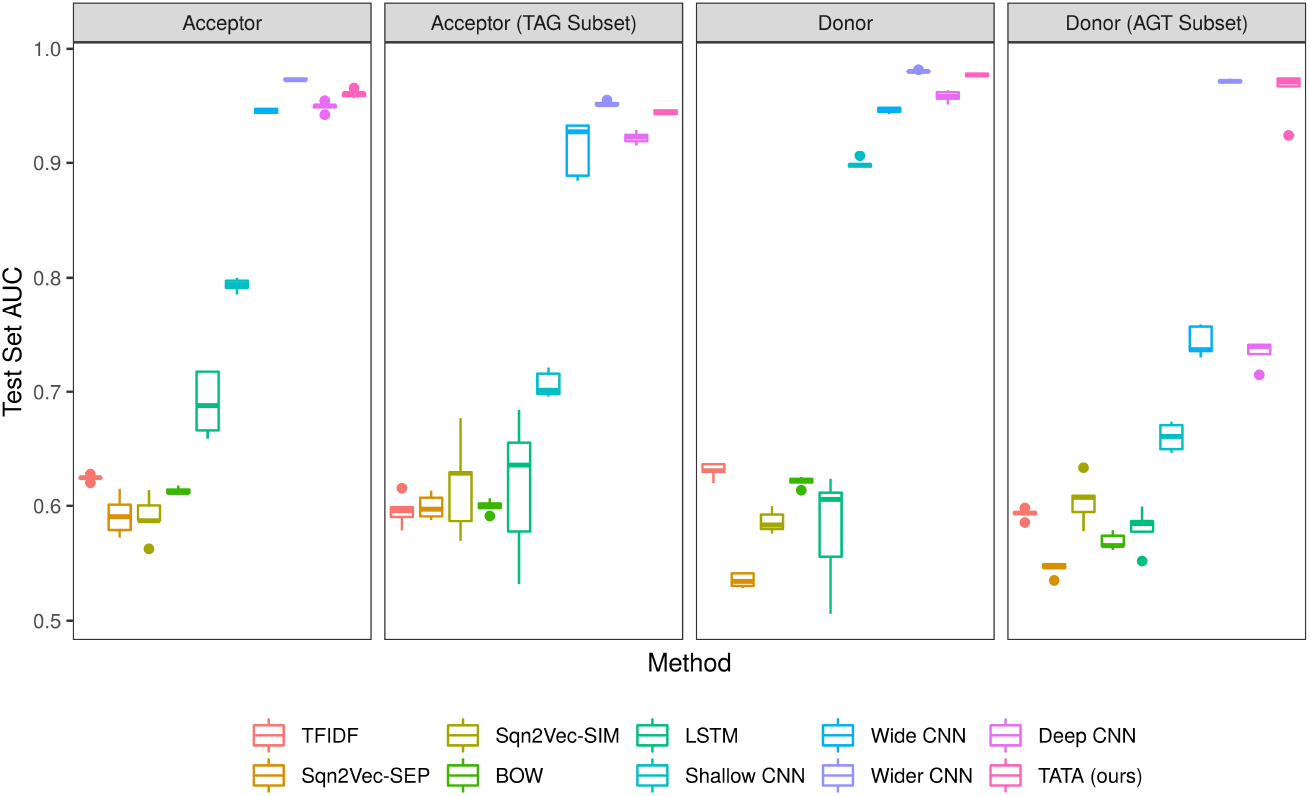
Test set AUC across 2 real data sets, using 5 independent training sets each. Our TATA architecture performs as well as, or better than, several state-of-the-art baselines. However, its parameters are explicitly interpretable. Here, we show TATA when *S* = 4 and *K* = 5.

In the case of the real splicing data, one could achieve decent accuracy by considering only the presence or absence of canonical splice motif, without considering context-dependent motif interactions. Non-canonical motifs occur less rarely, and are a less precise predictor of RNA splicing (i.e., a non-canonical motif may appear in a sequence without causing a splice may event). We hypothesize that non-canonical splice sites are harder to predict without also considering contextdependent motif interactions. Therefore, we subset the Albaradei et al. [2020] data to include only specific non-canonical splice sites in the positive class. Although we do not know the ground-truth mechanism, we can inspect the parameters of our neural network directly to generate hypotheses about how non-canonical splice sites might get recognized. For this, we viewed the visual-semantic representations of the model across 5 folds and *K* = {1, 2, 5} Toeplitz heads. Figure 7 presents a clear example of a pattern that appears more generally within the TAG-only acceptor subset data. Here a TA-rich sequence precedes a GC-rich sequence by about 50 base pairs. The poly-pyrimidine tract is defined by a repetition of Ts and As, and is known to mediate acceptor site splicing [Ohno et al., 2018]. Splice enhancers are often defined by Gs and Cs [Cartegni et al., 2003], and are known to play an important role in splicing when other canonical elements are lacking [Murray et al., 2008]. From this, we can formulate a testable hypothesis that having a GC-rich splice enhancer ~ 50 bases ahead of a poly-pyrimidine tract signals causes the RNA splicing machinery to recognize the non-canonical splice motif as a valid splice site.

**Figure 7:**
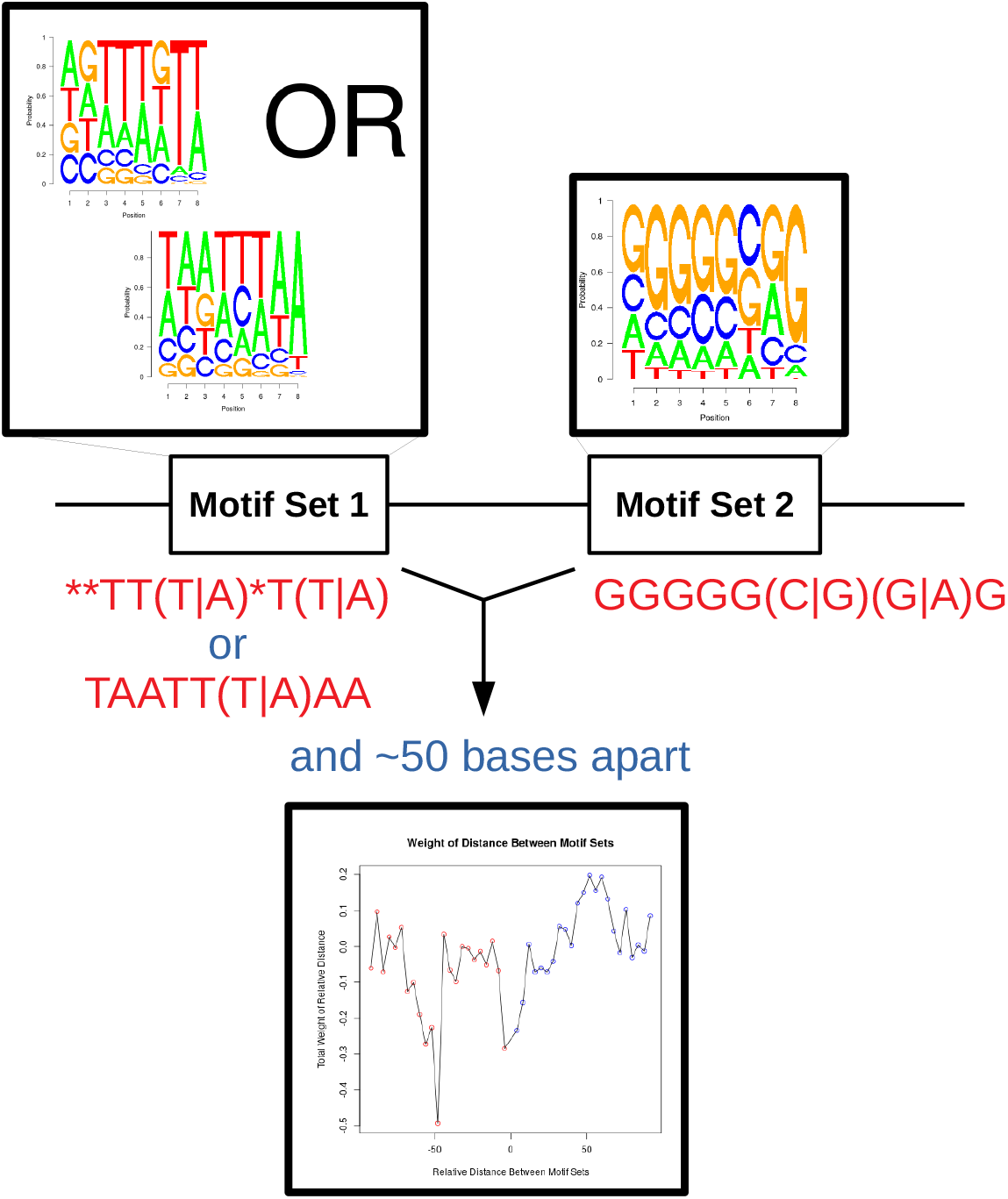
A concise visual-semantic representation of one Toeplitz Max Pool Head and all associated parameters. This representation, learned from the TAG-only acceptor subset data, can inform a testable hypothesis about a mechanism in non-canonical splice site detection: having a GC-rich splice enhancer ~ 50 bases ahead of a poly-pyrimidine tract signals causes the RNA splicing machinery to recognize the non-canonical splice motif as a valid splice site.

## 7 Conclusions

Here, we propose the TATA architecture for learning distance-dependent motif interactions through a novel Toeplitz max pool layer, built on top of a modified self-attention head. As a software tool, TATA can advance research into the basic biology of RNA splicing by generating a concise, human-interpretable knowledge representation of RNA splicing. This knowledge representation can help generate hypotheses about putative splice mechanisms, feeding back into the research cycle via a bit-to-bedside pipeline in which computational discoveries inform subsequent biomedical experiments. Although we focus on RNA splicing here, the method is generally applicable to other problems where distance-dependent motif interactions trigger genomic events.

## 8 Availability of Data and Materials

All data and code will become publicly available after peer review.

## Footnotes

1 The elements of **T** are always non-negative because **Q** and **K** are always non-negative. A ReLu activation is placed on **Ĥ** while a non-negative weights constraint is placed on {**W**_*Q*_, **W**_*k*_}.

2 The acceptor and donor data are artefactually centered on the labelled splice site. For these data, the final strided max pool layer is replaced with a global max pool layer to erase the positional information.

